# Correlated and Anticorrelated Binocular Disparity Modulate GABA+ and Glutamate/glutamine Concentrations in the Human Visual Cortex

**DOI:** 10.1101/2024.09.12.612491

**Authors:** Jacek Matuszewski, Ivan Alvarez, William T. Clarke, Andrew J. Parker, Holly Bridge, I. Betina Ip

**Affiliations:** Wellcome Centre for Integrative Neuroimaging, Nuffield Department of Clinical Neurosciences, University of Oxford, Oxford, United Kingdom, OX3 9DU; Laboratory of Brain Imaging, Nencki Institute of Experimental Biology, Polish Academy of Sciences, Warsaw, Pasteur 3, Poland; Institut für Biologie, Otto-von-Guericke Universität, Magdeburg 39120, Germany; Department of Physiology, Anatomy and Genetics, University of Oxford, Oxford OX1 3PT, United Kingdom

## Abstract

Binocular disparity is used for perception and action in three dimensions. Neurons in the primary visual cortex respond to binocular disparity in random dot patterns, even when the contrast is inverted between eyes (false depth cue). In contrast, neurons in the ventral stream largely cease to respond to false depth cues. This study evaluated whether GABAergic inhibition is involved in suppressing false depth cues in the human ventral visual cortex.

We compared GABAergic inhibition (GABA+) and glutamatergic excitation (Glx) during the viewing of correlated and anticorrelated binocular disparity in 18 participants using single voxel proton magnetic-resonance spectroscopy (MRS). Measurements were taken from the early visual cortex (EVC) and the lateral occipital cortex (LO). Three visual conditions were presented per voxel location: correlated binocular disparity; anticorrelated binocular disparity; or a blank grey screen with a fixation cross. To identify differences in neurochemistry, GABA+ or Glx levels were compared across viewing conditions.

In EVC, correlated disparity increased Glx over anticorrelated and rest conditions, also mirrored in the Glx/GABA+ ratio. In LO, anticorrelated disparity decreased GABA+ and increased Glx. Joint effects on GABA+ and Glx were summarised by the Glx/GABA+ ratio, which showed increased excitatory over inhibitory drive to anticorrelated disparity in LO. Glx during viewing of anticorrelation in LO was predictive of its object-selective BOLD-activity.

We provide evidence that early and ventral visual cortices change GABA+ and Glx concentrations during presentation of correlated and anticorrelated disparity, suggesting a contribution of cortical excitation and inhibition in disparity selectivity.

**Significance Statement:** The visual system must correctly match elements from the left and right eye for proper reconstruction of binocular depth. At the earliest part of binocular processing, false matches can activate depth detectors, however, the activation to false matches is absent in the ventral visual stream. We tested whether GABAergic inhibition contributes to the suppression of false matches in the ventral stream by measuring GABAergic inhibition and glutamatergic excitation in the human visual cortex during the presentation of correct and false matches. Correct matches increased excitation in response in the early visual cortex, and false matches increased excitation and decreased in the ventral visual cortex. These results suggest a role for excitation and inhibition in distinguishing depth cues for stereoscopic vision.

## Introduction

The act of seeing requires a transformation of retinal signals to perceptual reality. How the visual system solves the many ambiguities it encounters along the way is an enduring question in neuroscience. A prime example comes from the binocular visual system, where non-matching features in the two eyes create a challenge known as the ‘correspondence problem’, as elements from each eye need to be matched correctly in the presence of multiple possible false matches (Marr & Poggio, 1979). One experimental approach to investigate the ability of the binocular visual system to eliminate these false matches is to use random dot stereograms (RDS) that carry disparity cues but have opposite contrast in the two eyes (‘anticorrelated’ stimuli, Fig. 1d) (Cogan et al., 1993). In V1, disparity-sensitive neurons show an inverted, and attenuated, tuning to these anticorrelated compared to correlated RDS (Cumming & Parker, 1997). Since these anticorrelated random dot stimuli do not ordinarily result in depth perception (Cumming et al., 1998) (with some exceptions (Tanabe et al., 2008)), it is thought that visual areas that are sensitive to false matches do not directly play a role in depth perception.

**Figure 1.**
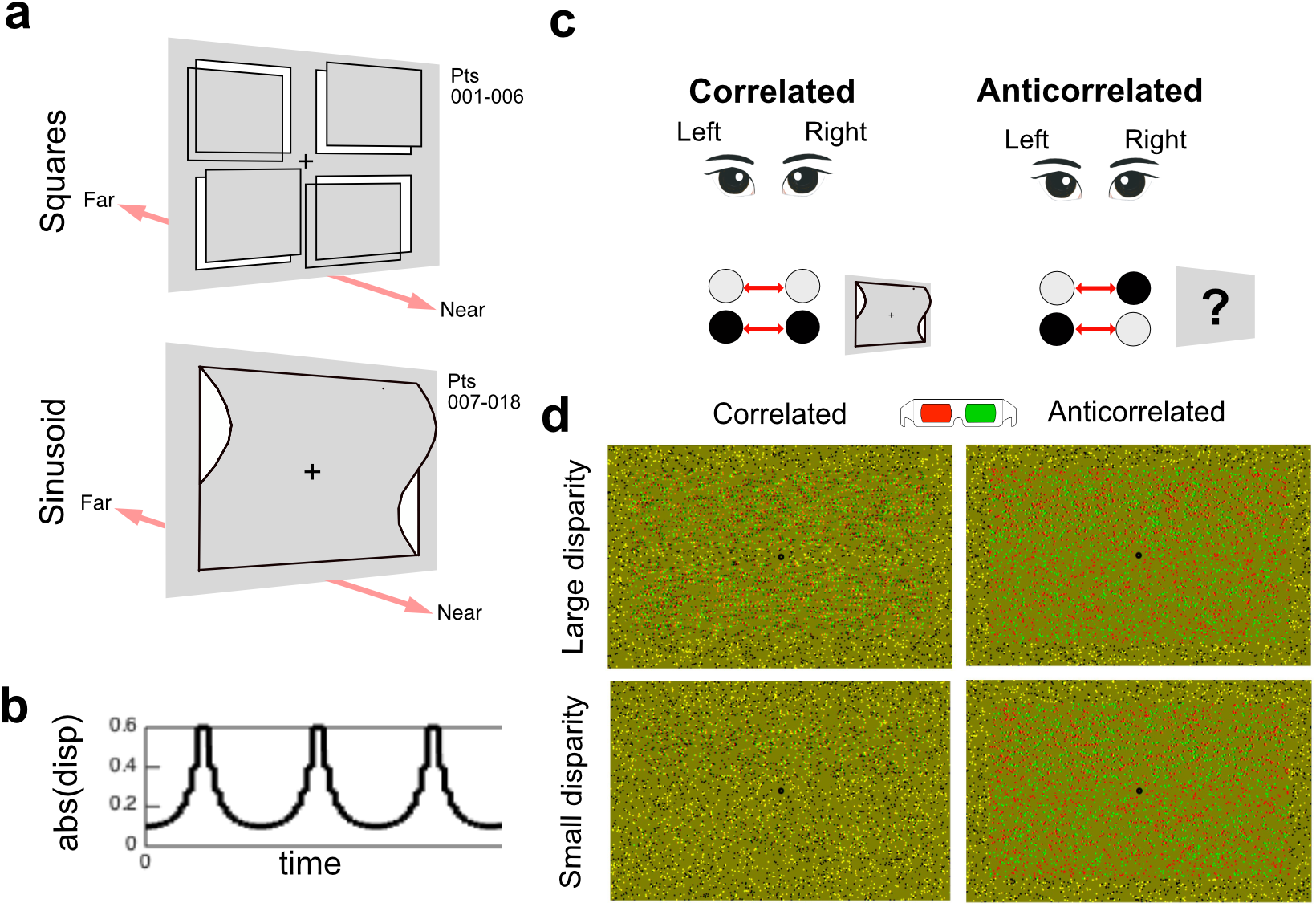
Diagram of random dot stereograms displayed inside the MRI-scanner. Two stimulus types were used which were consistent within participants: **(a)** Stimuli were composed of dynamic random dots stereograms (RDS) appearing like depth-defined squares for participants (Pts) 001-006, or spatial sinusoids, moving closer and farther in depth to the fixation plane over time, for participants (Pts) 007-018. Disparity type did not affect GABA+ concentrations, and data were pooled together. **(b)** example maximum absolute disparity over time. Participants were instructed to monitor the luminance of the stimuli and press a button on a button box when the luminance increased. The main conditions were **(c)** correlated disparity, which presented matching contrast to dots presented to the left and right eye, and anticorrelated disparity, which switched the contrast of the dots. White dots were matched with a black dot, and black dots with a white dot. Anticorrelated disparity did not evoke any depth percept. **(d)** Example images of sinusoid RDS in anaglyph, with large disparity amplitude **(top, left)** and small disparity amplitude **(bottom, left)**. Example anticorrelated images are to the right **(top, right and bottom right)**.

The stereo correspondence problem is solved in the ventral visual cortex of macaque (Janssen et al., 2003), yet the neural mechanisms by which false matches are suppressed and true matches are facilitated along the visual pathway remain poorly understood. The neuronal response to anticorrelated stimuli decreases along the ventral visual pathway (Kumano et al., 2008) until it ceases in the inferior temporal cortex (Janssen et al., 2003). Ventral visual cortex neurons only respond to true matches that are relevant for perception (Abdolrahmani et al., 2016; Janssen et al., 2000a; Yoshioka et al., 2021), a property that could support depth-invariant object recognition. Haemodynamic responses measured non-invasively from human participants largely support a loss of selectivity to anticorrelation in the lateral occipital area (LO) (Bridge & Parker, 2007; Preston et al., 2008), the putative homologue of macaque inferior temporal cortex. In contrast, dorsal extra striate areas MT (Krug et al., 2004) and MST (Takemura et al., 2001) of macaque retain sensitivity to anticorrelated stimuli. These findings suggest that false matches are partially and progressively removed in the ventral visual stream.

How is this suppression accomplished? Several potential and not mutually exclusive physiological mechanisms have been proposed that could account for the disparity selectivity observed (Read & Cumming, 2007). The two main theories (see review by (Verhoef et al., 2016)) are (i) feedback from more anterior areas in the visual ventral stream, such as area IT, that have a solution for the correspondence problem or (ii) refinement of selectivity through local inhibition in V1. A purely feedback-based mechanism is not supported by the literature. The time course of V4 neurons to anticorrelated disparity is not consistent with a feedback mechanism, as attenuation is instantaneous rather than delayed, as would be expected if a purely feedback-based mechanism was involved (Tanabe et al., 2004). Modifying the binocular energy model with feedforward local processing can account for the observed anticorrelated response in V1 (Read et al., 2002) and therefore support a role for cortical inhibition. Disparity tuning results from a combination of excitatory and inhibitory activity, as V1 neurons partially suppress false matches, caused by either inhibition or loss of excitation through prior inhibition (Tanabe & Cumming, 2014; Tanabe et al., 2011). Invoking a model with inhibition from local interactions additionally accounts for further anticorrelated disparity responses in V1 (Samonds et al., 2013). Adaptation to anticorrelated disparity increased the EEG measure of visual excitation, possibly signalling larger inhibitory activity for processing anticorrelated disparity in the visual cortex (Rideaux et al., 2020). Overall, these studies provide indirect evidence suggesting that GABAergic inhibition, which plays a role in neural suppression, may be involved in the processing of anticorrelated disparity. No study has directly probed GABAergic inhibition in relationship to anticorrelated disparity processing.

We measured GABAergic inhibition using the GABA+ signal from the early visual cortex (EVC) and the lateral occipital cortex (LO) in the human brain. Participants viewed correlated and anticorrelated binocular disparity during measurements. We also measured signals relating to glutamatergic excitation, using the glutamate+glutamine complex (Glx). Our results showed surprisingly that viewing of anticorrelated stimuli decreased inhibition and increased excitation in ventral area LO. A similar pattern was not present in the early visual cortex. These results are considered in relation to the stages of processing of binocular information in the human visual cortex.

## Methods

### Participants

Eighteen participants (thirteen females, mean = 29; SD = 8 years) took part in the experiment. All had normal, or corrected-to-normal, vision and demonstrated normal stereo acuity using clinical screening plates (<120 arcsecs, TNO Stereo test, Lameris, Utrecht). They had no self-identified neurological impairments. Before recruitment, all participants were verbally screened during the eligibility assessment for drug use and consumption of no more than three cigarettes per day. Each volunteer took part in a two-hour MRI-session. A reimbursement of £40 was received for the MRI-sessions. All volunteers gave informed and written consent, as approved by the local Research Ethics Committee (R54411/RE001). We did not perform an a priori power calculation for the study.

### Experimental set-up

A custom-made Wheatstone MRI-stereoscope was used for dichoptic presentation inside the MRI-scanner (Ip et al., 2022). Inside the scanner, participants verbally confirmed seeing stereoscopic depth during a brief calibration procedure prior to data collection. Participants viewed stimuli on a gamma-linearised BOLD screen (BOLDscreen 32, Cambridge Research Systems, viewing distance = 127.5 cm). The dichoptic display size in degrees of visual angle was 10.23° x 12.88°. All participants wore transparent protective goggles with or without MRI-safe visual correction.

### Visual stimuli

**Figure 1**. shows the experimental conditions. Two different disparity-defined stimuli were used but kept consistent within participant (**Fig 1a, top**): six participants were presented with a rectangular patch in which the peak disparity of four small quadrants varied between ±0.3° in logarithmic steps in depth without going through ± 0.05. The top right and bottom left quadrant were matched in disparity, and the top left and bottom right quadrant were matched in equal and opposite disparity. The stimulus was changed to a sinusoid to further optimise presentation for lateral occipital cortex. The remaining 12 participants were presented with a RDS stimulus appearing like a spatial sinusoid (**Fig 1a, bottom**), in which the peak disparity varied between ±0.6° in logarithmic steps in depth without going through ±0.1. In six out of these 12 participants, the stimulus briefly appeared completely flat before transitioning to ±0.1. For a detailed description of the stimuli see **Supplementary Materials, Table S3**. A linear mixed model showed that the stimulus type (‘square’, ‘sinusoid’) had no effect on spectral quality or GABA+ concentration and conditions were analysed together. RDS filled the entire display screen and consisted of 5,000 dots (50% white and 50% black, dot radius = 0.05°, dot refresh rate = 30 Hz). Stimuli subtended 7.23° x 9.88° and were framed by a 1.5° correlated, zero disparity border. A central fixation dot (black, 0.4°) was present on each frame, surrounded by a dot-free zone (0.8°) to prevent vergence eye movements caused by changing disparities at fixation. The disparity in the RDS modulated from positive to negative disparity over time to reduce adaptation effects by cycling six times over the MRS acquisition runtime (**Fig 1b**). Anticorrelated RDS were identical to correlated RDS (**Fig 1c**), with the exception that the interocular contrast of the dots was inverted and no perception of depth was reported (**Fig 1d**). During correlated and anticorrelated runs, participants looked at the fixation dot and performed an easy contrast detection task. Contrast differences occurred at random points during the run (72 times/run, 1s in duration) and comprised a 20% linear decrease in contrast across the stimulus area. The rest condition was a mid-grey screen with a fixation dot. Participants were asked to fixate, and no task was provided. We collected six MRS runs per participant (rest, correlated and anticorrelated conditions in both EVC and LO). Across participants half had EVC data collection first and the other half LO. Within each of the ROIs, the presentation order of rest, correlated and anticorrelated disparity was randomised. Each MRS acquisition lasted a total of 8 min and 13s, including dummy scans.

### MRI data acquisition

Magnetic Resonance images were acquired using a 3T Siemens Prisma (Siemens Healthineers AG, Erlangen, Germany), using a 64-channel head and neck coil. A 1 mm isotropic whole-head T1-weighted image was collected for placement of spectroscopic volume of interest (MPRAGE, TR=1900 ms; TE=3.97 ms; field-of-view= 192×192 mm^2^; 192 slices; flip angle=8°), with a total acquisition time of 5 min 31s. To locate the lateral occipital cortex region, an object-localiser scan was performed prior to MRS voxel placement (2.4 mm isotropic resolution; MB8; TR=1000 ms; TE=39 ms; 66 slices; flip angle = 52°). Stimuli were presented in a block design, with intact objects (15s) alternating with phase scrambled objects (15s) that preserved the same contrast and phase information as intact objects, while removing shape information. The total acquisition time was 3 min 10s. A fixation cross was always present, and each trial was presented for 1000 ms (800 ms stimulus followed by 200 ms mid-grey screen). Participants were instructed to passively fixate during the localiser experiment.

### MRS procedure

Single-voxel MRS data were acquired using a MEGA-PRESS sequence, consisting of a locally developed version of the CMRR spectroscopy package MEGA-PRESS sequence (Willis et al., 2023). Acquisition parameters consisted of: voxel size = 20×25×25 mm^3^ (short dimension in the medial-lateral direction for LO, and with the long dimension aligned with the tentorium cerebelli for EVC); echo time (TE) = 68 ms; repetition time (TR) = 1500 ms; number of spectra = 320 (160 edit-on and 160 edit-off spectra); VAPOR and dual-band editing pulse water suppression; 22.3 ms editing pulse for a 53 Hz bandwidth, which was centred at 1.9 ppm (edit-on condition) and at 7.5 ppm (edit-off condition) in alternation; 16-step phase cycling; a total of 8 min 13 s run time. A water unsuppressed reference was collected after each condition to provide an internal reference.

The region of interest for EVC was first centred to the occipital midline to cover equivalent portions of the right and left visual cortex, then angled to be parallel to the calcarine sulcus and moved as posterior as possible while avoiding contamination by the cerebellar tentorium and the sagittal sinus (**Fig 2a**). For LO placement, the voxel was positioned in the right lateral occipital cortex using the object localiser. Data were analysed on-line using dynamic t-maps of the contrast between objects and phase-scrambled objects. The right hemisphere has been shown to respond more to disparity processing than the left (Ip et al., 2014; Lohia et al., 2024). Contrasting the intact with phase-scrambled objects isolated regions where haemodynamic responses increased to intact objects, as shown by activation clusters in bilateral ventral visual cortex (**Fig 2b**). In confirmation of the accuracy of the voxel placement, we found substantial %BOLD-change to objects inside the LO voxel (**Fig 2c, top**, n = 18, mean = 0.68 ± 0.23), and 0.66 proportion overlap between the LO voxel with the statistical maps from the object localiser (**Fig. 2c, bottom**, n = 18, mean = 0.66 ± 0.10).

**Figure 2.**
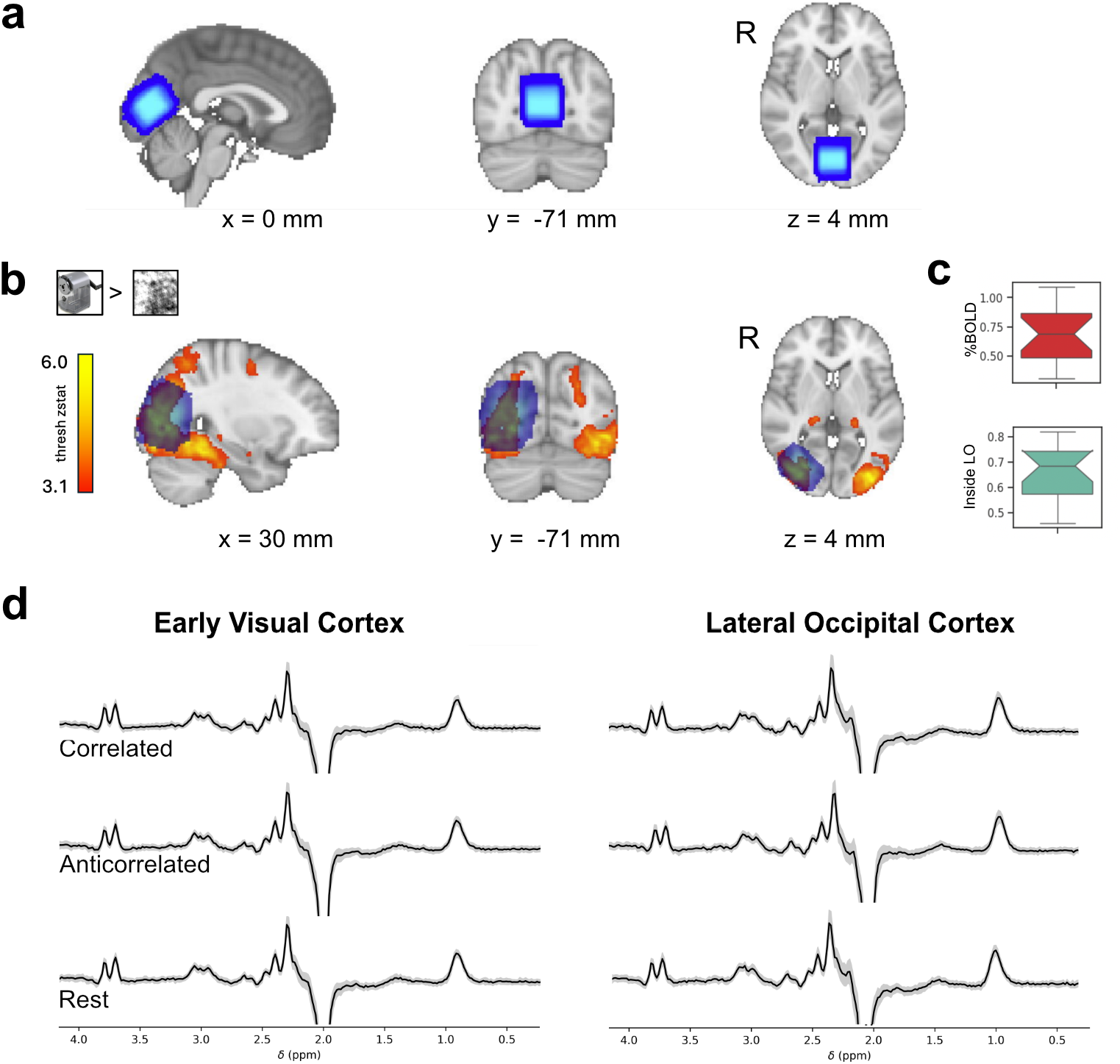
MRS voxel in early and ventral visual cortex and group spectra for each condition. **(a)** shows proton magnetic resonance spectroscopy (MRS) voxel locations for all 18 participants positioned in the early visual cortex (EVC; blue) and **(b)** right lateral occipital cortex (LO; blue) displayed on the MNI-152 2 mm standard. **(c)** boxplots show %BOLD-change to objects inside the LO voxel, and proportion overlap between statistical activation maps to objects and the LO voxel. **(d)** Group MRS spectra for each condition and voxel. The black line represents the mean across participants, the grey area shows standard deviations across participants. Spectral amplitude is scaled to arbitrary units. X-axis shows chemical shift in parts per million. Slice positions are indicated in MNI-coordinates. The boxplot shows the middle 50% of points, and the upper and lower whiskers span points outside the middle 50%. R = right (hemisphere).

### fMRI analysis

For each subject, functional data from the object localiser were pre-processed and analysed with FSL FEAT (version 6.0.7.10, http://www.fmrib.ox.ac.uk/fsl). This pipeline included realignment and head movement correction (MCFLIRT), registration to standard MNI152 space (2 mm resolution), high pass filtering (30 s cut-off) and spatial smoothing with 5 mm Gaussian full-width half maximum (FWHM) kernel. Next, timings of intact and phase scrambled object conditions and six standard head motion parameters were entered into subject-specific first-level general linear models (GLM). Finally, *object > scrambled* contrasts from all subjects were entered into a group-level mixed effects analysis (FSL FLAME 1). Group-level maps were cluster-corrected for multiple comparisons with a Z threshold of 3.1 (*p*< 0.05) and overlapped with the group mask of LO MRS voxel created by joining all subjects’ masks. Additionally, to investigate BOLD signal responses in the LO MRS voxel location, we computed intersections between each subject’s functional maps and LO MRS voxel with `fslmaths`. Then, we used `featquery` to calculate the mean BOLD-change (%) in the obtained mask.

### MRS data analysis

Data analysis was performed using the open-source toolbox FSL-MRS v.2.1.19 (Clarke et al., 2021). Processing included conversion from the vendor .dat (or TWIX) format to NIfTI format using spec2nii v.0.7.4 (Clarke et al., 2022), pre-processing included: coil-combination, windowed averaged phase and frequency alignment between repeats, eddy current correction, truncation of the FID to remove two time-domain points before the echo center, removal of residual water peak using Hankel Lanczos singular value decomposition (HLSVD) over 4.5 – 4.8 ppm, phase and frequency alignment between averaged edit-on and edit-off spectra. A phase-corrected, non-water-suppressed reference was also generated during pre-processing from integrated reference scans. Model-fitting was performed using a Linear Combination model: basis spectra were fitted to the complex-valued spectrum in the frequency domain by scaling, shifting, and broadening them. Then, basis spectra were divided into 2 metabolite groups, with macromolecular peaks allowed to broaden and shift independently of other metabolites. The model fitting was achieved using the truncated Newton algorithm as implemented in Scipy. A 0^th^-order complex polynomial baseline was fitted concurrently. To model metabolites in the edit-on minus edit-off difference spectrum, we used a simulated basis set containing the model spectra for N-acetylaspartate (NAA), N-acetylaspartateglutamate (NAAG), γ-amino-butyric acid (GABA), glutamine (Gln), glutamate (Glu), glutathione (GSH), macromolecules (MM) and combined NAA+NAAG, Glu+Gln+GSH, GABA+MM (https://git.fmrib.ox.ac.uk/wclarke/win-mrs-basis-sets). As the GABA signal at 3.0 ppm contains co-edited macromolecule signals, as well as homocarnosine (Rothman et al., 1997), the combined GABA+MM signal is reported and referred to as GABA+. The combination of Glu+Gln+GSH is referred to as Glx. Both Glx and GABA were extracted from the difference spectrums. Absolute concentrations in millimole per kilogram (mMol/kg), corrected for tissue fraction and tissue relaxation are reported. FSL FAST was applied to the T1-weighted image for voxel tissue type fraction estimation, separating white matter, grey matter, and cerebrospinal fluid, for tissue fraction correction.

### MRS data quality measure

Prior to summary statistics, an outlier analysis based on metabolite concentrations was performed to ensure that data represented the norm. GABA+ and Glx concentrations outside of the 1.5 times the interquartile range were excluded. Some conditions could not be collected from all participants, and MRS quality control excluded some individual data points. We ended up with a minimum of 15 participants per condition. For a full breakdown of data points, see Supplementary Materials, **Table S1**. The spectral quality of the acquisition was determined using the linewidth of the full-width at half-maximum (FWHM) of the inverted N-acetylaspartate singlet at 2.01 ppm and reported in Hz. The GABA+ model fit was reported as absolute Cramer-Rao Lower Bounds (CRLB, **Table S2)**. We assessed MRS data quality across voxel locations and conditions. There was no interaction effect of voxel and condition on NAA linewidth (*F*_1,77.275_ = 0.42, *p* = 0.66), meaning that acquisition quality did not depend on visual condition. A model without interaction showed a highly significant effect of voxel on spectral quality (**Fig. 3a**, *F*_1,79.12_ = 183.73, *p* <2e-16), with LO linewidth broader than EVC. No effects of condition were found (*F*_1,79.20_ = 0.079, *p*=0.92). In contrast, no effect of voxel or condition were found on the quality of the GABA+ model fit (**Fig. 3b**, *p* > 0.05). Based on the linewidth difference, we performed separate analyses for EVC and LO voxels as they could not be directly compared and added NAA linewidth as a covariate of no interest in the LMM analyses. We also controlled for linewidth in the partial correlations.

**Figure 3.**
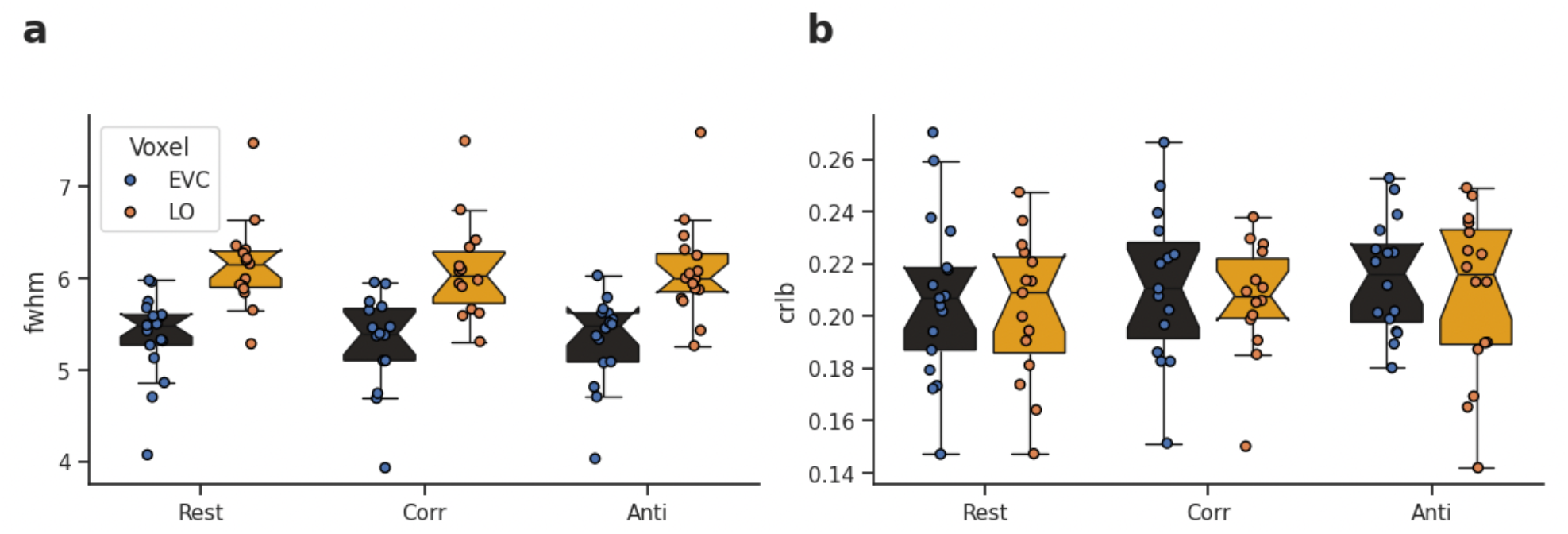
MRS quality measures in early visual cortex (EVC) and lateral occipital cortex (LO). Boxplots show spectral quality measure NAA linewidth by voxel and condition **(a)**, and goodness of model fit measure absolute Cramer-Rao Lower Bounds (CRLB) for GABA+ by voxel and condition **(b)**. Black boxplot, blue dots = EVC, orange boxplot, orange dots = LO. The boxplot shows the middle 50% of points, and the upper and lower whiskers span points outside the middle 50%. Corr = correlated condition; Anti = Anticorrelated condition; FWHM = peak full width at half maximum.

### Statistical analysis

Data analysis was conducted using RStudio (RStudio Version 2023.06.0+421). Metabolite concentrations were analysed with mixed effects linear models computed with LME4 R package (Bates D, 2015). The model syntax was constructed to test for fixed effects of condition within voxel, including the dummy variable ‘spectral quality’, and the random effect of ‘participant’. Separate analyses were performed for each voxel (EVC, LO), and each metabolite (GABA+, Glx). A model without interaction was used if no interaction effects were found. The anova function was used to obtain *p* values. A Type II Analysis of Variance Table with Kenward-Roger’s method was used when no interactions were present. Each linear model was performed with an expression: *metabolite_concentration ∼ Condition + NAA FWHM + (1*|*Participant)*. If a fixed effect was significant, posthoc analyses were performed using Tukey method for multiple comparisons of means for parametric models with p-values adjusted using Bonferroni-Holm correction (R, multcompare): *summary(glht(model, linfct = mcp(cond = “Tukey”)), test = adjusted(“holm”))*. Pearson’s correlations coefficients with and without partial correlations were computed with pingouin (Vallat, 2018) to assess the relationship between GABA+ and Glx in each voxel and condition. The excitation/inhibition index was calculated by dividing Glx/GABA+ using the same data points as for the individual metabolites analyses. We applied two-tailed Fisher’s *r*-to-*z*-transform tests (http://vassarstats.net/rdiff.html) to test the significance of the difference in correlation coefficients.

## Results

We evaluated if GABA+ in the early or ventral visual cortex was modulated by presenting correlated or anticorrelated random dot stereograms, with identical disparity magnitudes but with inverted contrast between eyes. As a baseline, we also presented a rest condition that required fixation on a mid-grey screen. Our hypothesis was that ventral visual cortex GABA levels would increase to anticorrelated disparity, whereas no change was expected in the EVC. We expected an increase in Glx during correlated and anticorrelated conditions compared to the rest, as a positive control for functional changes in glutamatergic excitation during visual stimulation. The summary metric Glx/GABA+ was calculated to evaluate changes in excitation and inhibition balance. In exploratory analyses, the relationship between LO GABA+ and Glx and the haemodynamic response to objects was assessed. Finally, metabolite concentrations from the early and ventral visual cortex allowed us to investigate the balance between GABA+ and Glx at two different stages of the visual processing hierarchy.

### Disparity correlation modulates GABAergic inhibition (GABA+) and glutamatergic excitation (Glx) in the early and ventral visual cortex

Viewing different stimulus conditions had no effect on GABA+ concentration in the EVC (**Fig.4a**, EVC, *p* = 0.64). When investigating the marker for glutamatergic excitation (Glx), we found that there was a significant effect of viewing conditions in the EVC (**Fig.4b**, *F*_2,29.38_ = 5.267, *p*=0.011). Specifically, seeing the correlated stimulus marginally increased Glx over anticorrelated, *p* = 0.071, and over Rest, *p*= 0.004. The excitation/inhibition-ratio (‘E/I-ratio’) evaluated the overall excitatory drive by dividing Glx by GABA+ (**Fig.4c**). Viewing conditions marginally modulated E/I-ratio in EVC (*F*_2,30.18_ = 2.87, *p*=0.072). Specifically, viewing correlated stimuli increased E/I ratio in the EVC relative to rest (*p*=0.067).

**Figure 4.**
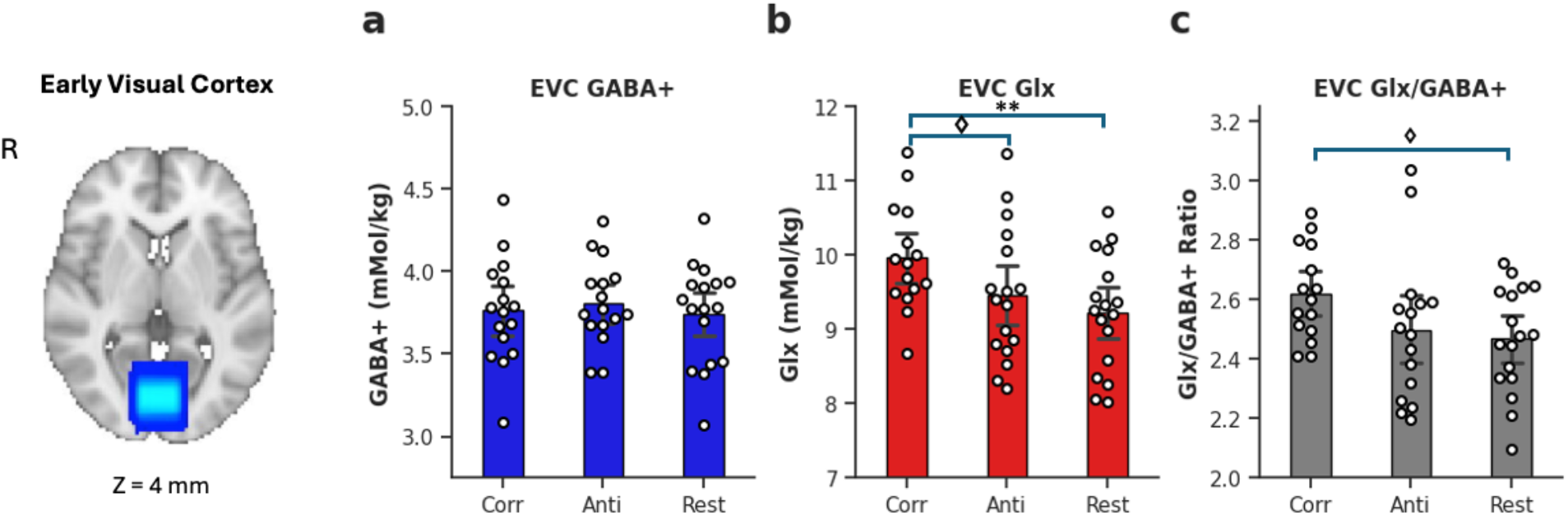
Bar plots show concentrations of metabolites for visual conditions for the early visual cortex. GABA+ **(a)**, Glx **(b**) and Glx/GABA+ (Excitation/Inhibition)-ratio **(c)**. Dots show individual participants. Error bars are +/-1 SD. Bonferroni-Holm adjusted p-values = * <0.05, ** <0.01, marginal significance ⋄ <0.1. Voxels show overlap across 18 participants. EVC = early visual cortex, GABA+ = γ-amino-butyric acid + macromolecules, Glx = glutamate + glutamine, Corr = correlated condition, Anti = anticorrelated condition. Slice positions are indicated in MNI-coordinates.

Viewing conditions modulated GABA+ in LO (**Fig.5a**, *F*_2,30.58_ = 3.42, *p*=0.047); viewing anticorrelation lowered GABA+ compared to rest (*p*=0.027). This was contrary to our hypothesis that GABA would increase during presentation of disparity anticorrelation. In LO, viewing anticorrelation marginally increased Glx compared to rest (**Fig. 5b**, *p*=0.055). Viewing condition influenced the E/I-balance in LO (*F*_2,29.1_ = 6.21, *p*=0.006); viewing anticorrelation increased E/I balance over correlation (*p*=0.018) and over rest (*p*=0.002).

**Figure 5.**
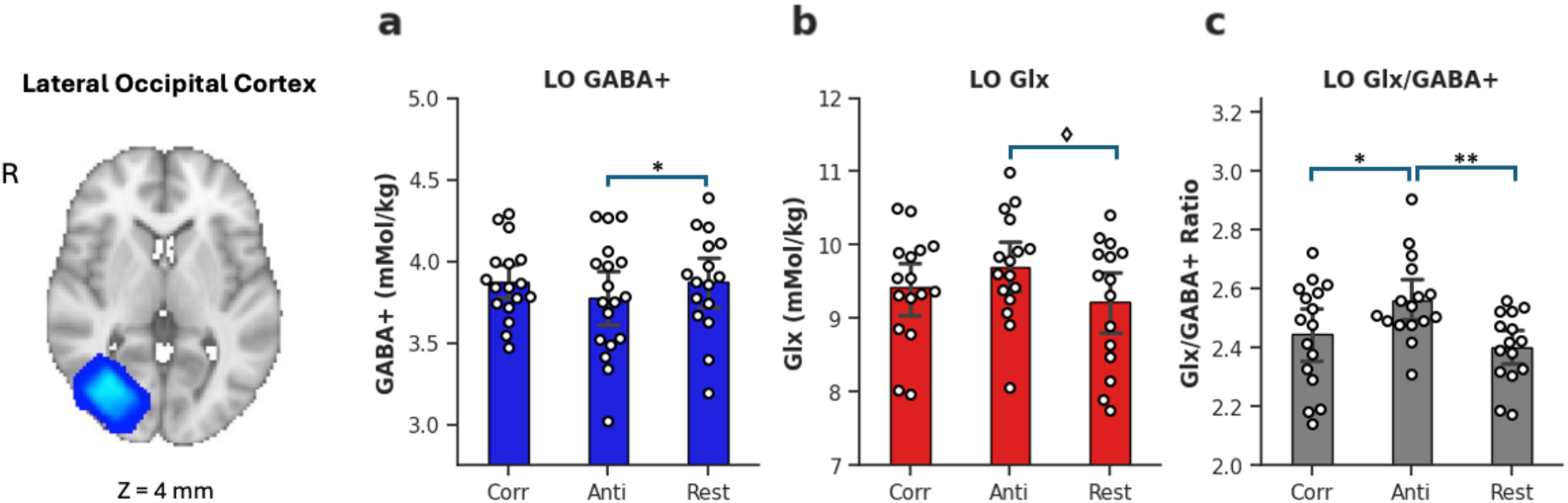
Bar plots show concentrations of metabolites for visual condition for the lateral occipital cortex. GABA+ **(a)**, Glx **(b)** and Glx/GABA+ (Excitation/Inhibition)-ratio **(c)**. Dots show individual participants. Errorbars are +/-1 SD. Bonferroni-Holm adjusted p-values = * <0.05, ** <0.01, marginal significance ⋄ <0.1. Voxels show overlap across 18 participants. LO = lateral occipital cortex; GABA+ = γ-amino-butyric acid + macromolecules, Glx = glutamate + glutamine, Corr = correlated condition, Anti = anticorrelated condition. Slice positions are indicated in MNI-coordinates.

In this section, we demonstrated that correlated binocular disparity increased cortical excitation in the early visual cortex, whereas anticorrelated disparity increased excitation in the ventral visual area LO. Excitatory drive in LO increased during anticorrelated disparity compared to correlated disparity or rest.

### Glutamatergic excitation (Glx) and GABAergic inhibition (GABA+) correlate in early and ventral visual cortex

The ratio of Glx/GABA is a proxy for excitatory and inhibitory balance for non-invasive brain imaging. The balance between excitation and inhibition is thought to enhance neuronal selectivity and support plasticity. In the human brain, their correlation has been shown to vary by brain region (Rideaux 2021, Steel 2020). While E/I balance has been studied in the early visual cortex, little is known about E/I-balance in the ventral visual cortex. Here we aimed to assess if their balance differed by disparity (defined by viewing condition), and by voxel location.

In EVC, we found that Glx and GABA+ were moderately correlated (**Fig 6a**, Rest, *r* = 0.63, *BF*_*10*_ = 8.44, *p* = 0.007; Corr, *r* = 0.65, *BF*_*10*_ = 7.12, *p* = 0.009). Only the anticorrelated condition did not show a significant correlation (*r* = 0.35, *BF*_*10*_ = 0.66, *p* = 0.206). In LO, we found weak evidence for a correlation between Glx and GABA+ in the correlated condition (*r*= 0.49, *BF*_*10*_ = 1.39, p = 0.076), but strong evidence (**Fig. 6b**, Anti, *r* = 0.72, *BF*_*10*_ = 15.47, *p* < 0.001; Rest, *r* = 0.85, *BF*_*10*_ = 403.63, *p* < 0.001) for this correlation for the anticorrelated and rest conditions respectively. Overall, the trend across conditions was positive, so data were pooled across conditions to investigate the E/I-balance across the two voxel locations. The correlation was extremely high in both EVC (**Fig. 6c**, *n*=48, *r*= 0.53, *BF*_*1*0_= 230.227, *p*=0.0001) and LO (**Fig. 6d**, *n*=47, *r*=0.65, *BF*_*10*_=6.83+04, *p*< 0.0001). These correlations were robust to controlling for several confounding factors separately, including spectral quality, participant age, sex, GABA+ model fit in EVC and LO. However, the correlation in EVC was not robust to controlling for Glx model fit, whereas it remained robust in LO (**Table 1**). When controlling for all confounding factors in the same model, the correlation between Glx and GABA+ was no longer significant for EVC (*r* = 0.214, *p* = 0.169), and the correlation in LO remained significant (*r* = 0.392, *p* = 0.0124). We assessed the significance of the difference in correlation coefficient between voxels using a Fisher’s r-to-z transform and found none (*p* = 0.358).

**Figure 6:**
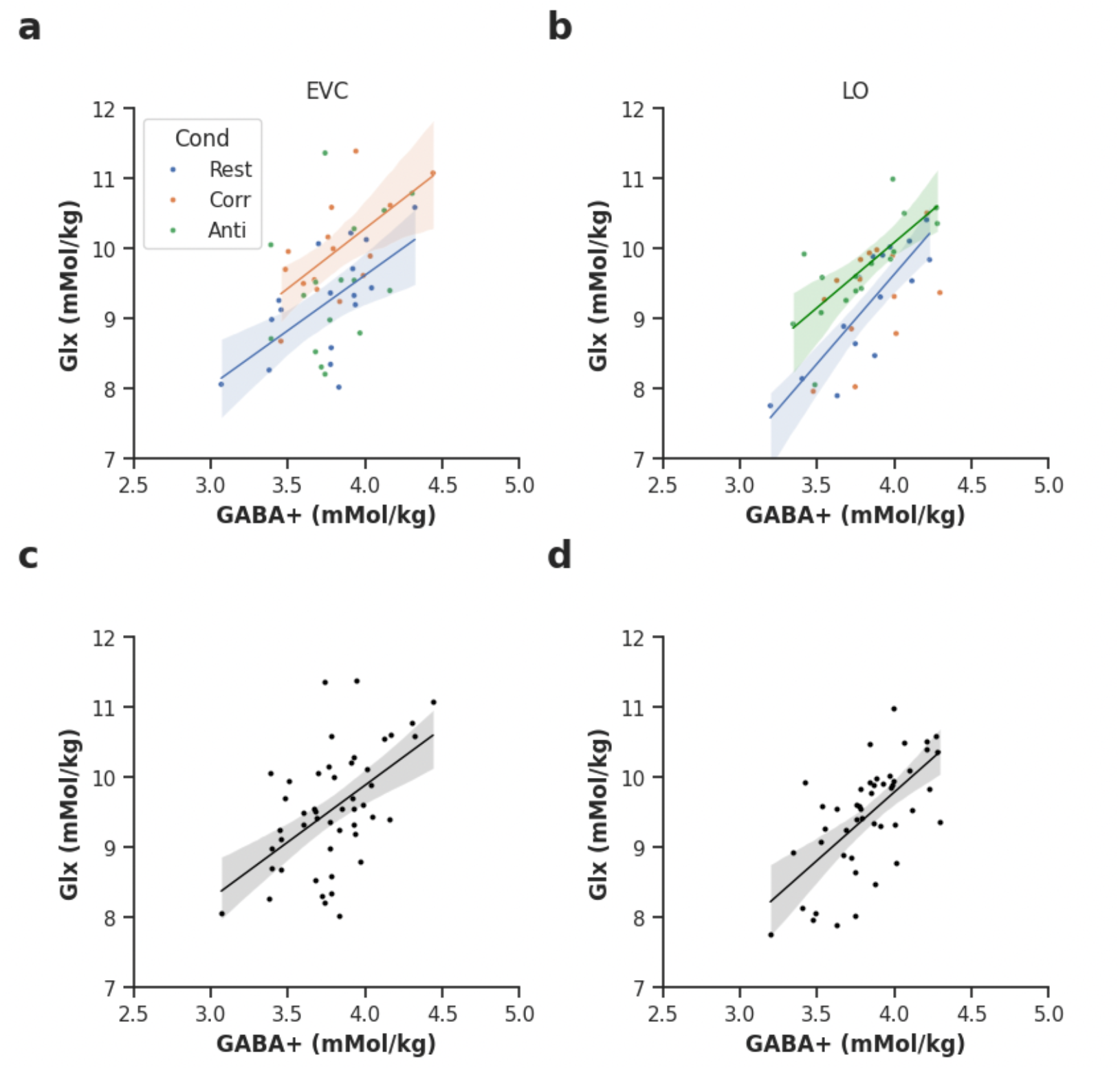
Glx and GABA+ were correlated within conditions for EVC and LO. Correlation between Glx and GABA+ for EVC **(a)** and for LO **(b)**. Pooled across conditions for EVC **(c)** and LO **(d)**. Model fit is plotted where p<0.05 uncorrected. Colours indicate condition, red = correlated (Corr), blue = rest, green = anticorrelated (Anti), black = pooled across conditions. GABA+ = γ-amino-butyric acid + macromolecules, Glx = glutamate + glutamine.

**Table 1:** partial correlation while controlling for confounding factors for the early visual cortex (EVC) and lateral occipital cortex (LO). P-values were Bonferroni-corrected for five multiple comparisons except for the full model. GABA+ = γ-amino-butyric acid + macromolecules, Glx = glutamate + glutamine + glutathione. * = p<0.05, **=p<0.01, ***=p<0.001, ****= p<0.0001, NS = not significant.

| Controlling for | *EVC*<br>*r* | *Corrected p-value* | *LO*<br>*r* | *Corrected p-value* |
| --- | --- | --- | --- | --- |
| Spectral quality | 0.53 | 0.0005 *** | 0.54 | 0.0005 *** |
| Age | 0.51 | 0.0015 ** | 0.63 | < 0.0001 **** |
| Sex | 0.48 | 0.003 ** | 0.65 | < 0.0001 **** |
| GABA+ fit error | 0.39 | 0.034 * | 0.45 | 0.00195 ** |
| Glx fit error | 0.30 | 0.209 NS | 0.60 | < 0.0001 **** |
| Spectral quality, Age, Sex, GABA+ fit error and Glx fit error | **0.214** | **0.169** | **0.392** | **0.0124 \*** |

These results show a tight balance of excitation and inhibition in the ventral visual cortex that was not driven by confounding factors.

### The correlation between object-selective BOLD-responses and glutamatergic excitation in the ventral visual cortex

Neurons in the ventral visual cortex exhibit two-dimensional (Decramer et al., 2019; Kourtzi & Kanwisher, 2000) as well as three-dimensional (Decramer et al., 2019; Janssen et al., 2000b) object selectivity. The relationship between 2D and 3D object processing in LO is not well understood. In this section, we evaluated the relationship between the excitation and inhibition during binocular disparity presentation in LO and 2D object selectivity. Using data from the LO voxel, we correlated the object-selective BOLD-responses with neurochemistry (**Fig. 1c**). Strong evidence was found for a correlation between object-selective BOLD-change and Glx only during viewing of anticorrelated disparity (**Fig. 7c**, *n* = 16, *r* = 0.7, *BF*_*1*0_= 18.77, *p* = 0.003). The correlation between LO anticorrelated Glx and BOLD-change was robust to controlling for spectral quality (*r* = 0.60, *p*= 0.017). To rule out the possibility that the BOLD-activity could have been higher in regions where there was more overlap between the LO voxel with the BOLD-activity, we performed partial correlations. The correlation remained significant after controlling for percentage overlap, (*r* = 0.67, *p* = 0.006). To evaluate if general excitability influenced the correlation, we controlled for EVC anticorrelated Glx concentrations and found no weakening of the relationship, *r* = 0.76, *p* = 0.002. The uncorrected correlation was significant after Bonferroni-correction for six multiple comparisons (*p* = 0.018). No comparable association was present for EVC Anti (*r* = 0.31, *p* = 0.24, *BF*_10_=0.575).

**Figure 7.**
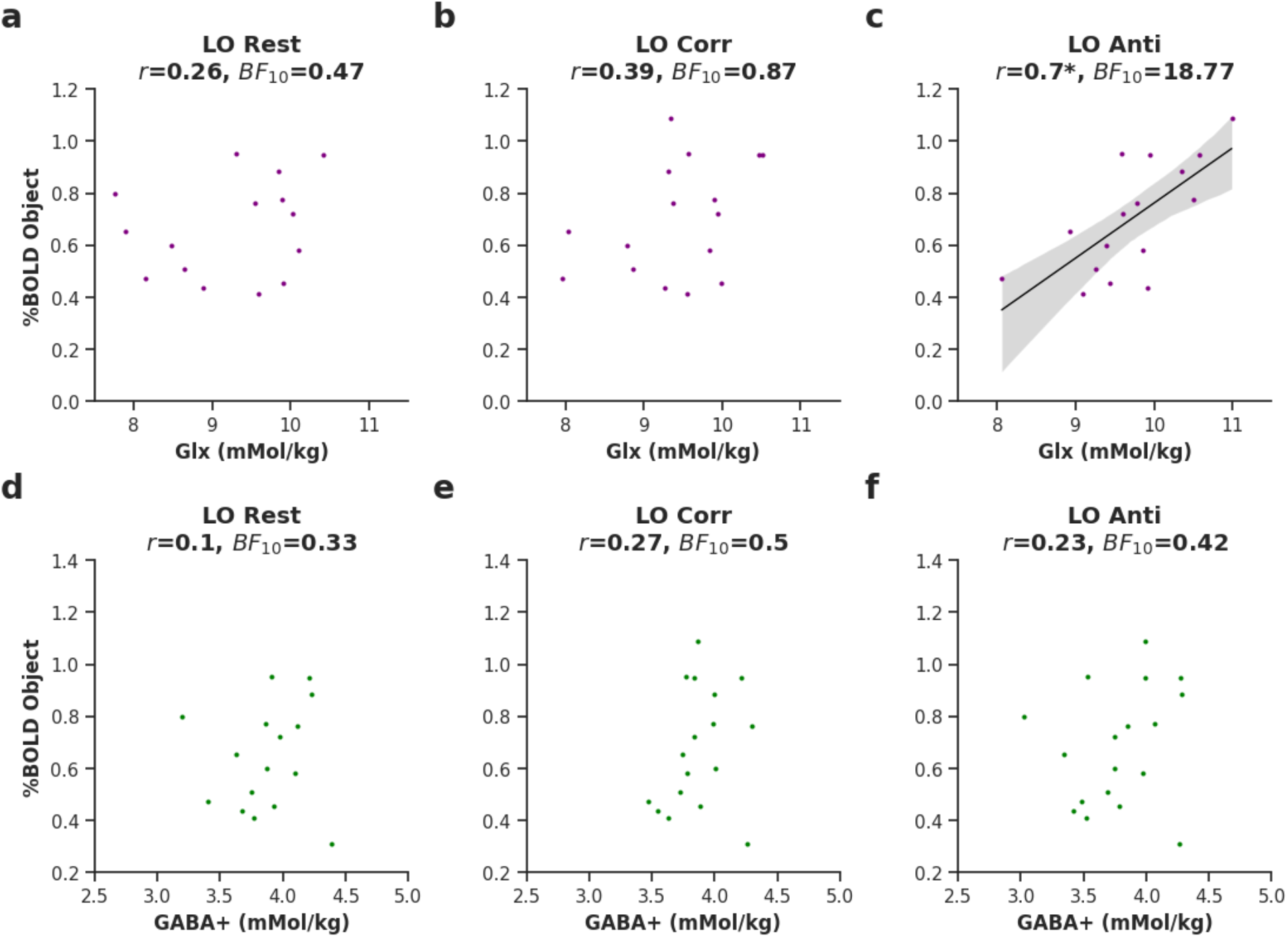
Correlation between %BOLD-change to objects with Glx and GABA+ in the lateral occipital voxel. Plots show correlation between %BOLD-change to objects with Glx during **(a)** rest, **(b)** correlated (Corr) and **(c)** anticorrelated (Anti) disparity in LO. Below, plots show the correlation between %BOLD-change to objects with GABA+ during **(d)** rest, **(e)** correlated, **(f)** anticorrelated disparity in LO. Model fit is plotted where p<0.05, Bonferroni-corrected. LO = lateral occipital cortex; GABA+ = γ-amino-butyric acid + macromolecules, Glx = glutamate + glutamine + glutathione, BF = Bayes Factor.

In summary, this section supplied strong evidence for a correlation between object-selective BOLD-activity and glutamatergic excitation during viewing of anticorrelated disparity, with greater BOLD-activity relating to more excitation. The correlation was robust to controlling for separately for spectral quality, percentage overlap and Glx in the early visual cortex.

## Discussion

Using proton MR Spectroscopy in humans, we measured GABAergic inhibition and glutamatergic excitation within the early and ventral visual cortex during viewing binocular disparity-containing RDS patterns. We show that switching the disparity correlation from correlated to anticorrelated, which removed the visual percept of depth, changed the neurochemistry in these areas: anticorrelated disparity decreased GABA+ and increased Glx in LO, resulting in an increased excitatory drive to anticorrelated disparity. By comparison, in the EVC, anticorrelated disparity decreased Glx. Ultimately, these results suggest that excitation/inhibition balance contributes to solving the stereo correspondence problem along the visual pathway.

### Increases in Glx to correlated disparity in early visual cortex

The primary visual cortex has been the intense focus of research on binocular correspondence. V1 neurons are strongly selective to binocular disparity and remain excited by anticorrelation (Cumming & Parker, 1997). However, the anticorrelated response in V1 is reduced and inverted, pointing towards an early (partial) suppression of false matches. Computational models propose that inhibition at this level suppresses anticorrelation (Read et al., 2002), a process that would reduce feedforward response to false matches at the earliest stage. While we found no change in GABA+, we did find a selective decrease in Glx to anticorrelated compared to correlated stimuli. Anticorrelated stimuli differ from correlated in several ways: firstly, the low-level matching is different, reflected in V1 neuronal responses; secondly, the correlated stimulus leads to a compelling percept of depth which may have top-down effects on responses; thirdly, anticorrelated stimuli can look like there are more dots, as not all are successfully matched (unpublished observation). An increase in Glx during correlated compared to anticorrelated disparity is consistent with a greater neuronal response to correlated disparity (Cumming & Parker, 1997; Janssen et al., 2003). Stereoscopic depth can also engage attentional modulation (Rose et al., 2003; Zou et al., 2022), potentially adding to the net excitation. In summary, the increase in excitation to disparity correlation compared to anticorrelation is consistent with a greater neuronal response to correlated disparity compared to the anticorrelated counterpart that lacks a coherent perceptual solution. The increase in excitation to correlation compared to rest is consistent with findings that show an increase in glutamate with visual stimulation (Bednarik et al., 2015; Ip et al., 2017; Kurcyus et al., 2018; Mangia et al., 2006). Overall, these results provide evidence that glutamatergic rather than GABAergic mechanisms may play a role in filtering false matches in the early visual cortex.

### Increases in Glx and decrease in GABA+ to anticorrelation ventral visual cortex

If binocular correspondence in natural vision is already complete before signals arrive in LO, then at the level of LO, signals from false matches would be absent in correlated RDS patterns. Alternatively, if GABAergic inhibition helped filter out false matches in LO, then we might expect to see an increase in GABA+, consistent with the idea of local suppression. Both outcomes would be consistent with strongly suppressed neuronal firing to anticorrelated disparity. Surprisingly, our results showed neither. We found a decrease in inhibition, together with an increase in excitation relative to correlated and rest conditions. This unexpected pattern might suggest that overall, anticorrelation broadly excited neurons in LO. Prior studies have found that anticorrelated stimuli caused less informative (Preston et al., 2008); and decreased hemodynamic response in LO compared to correlated stimuli (Alvarez et al., 2021; Bridge & Parker, 2007; Ip et al., 2022). These population-level measurements contrast with neurophysiological findings that show a complete cessation of response in single neurons (Janssen et al., 2003).

One possibility is that adaptation to anticorrelation over the presentation time of the stimulus caused the balance to shift towards excitation. In support, a prior study has shown an increase in the EEG measure of excitability during presentation of anticorrelated and not correlated disparity (Rideaux et al., 2020). Excitation during the rebound phase of adaptation could thus signal a primary suppressive response to binocular anticorrelation, consistent with the idea that GABAergic inhibition suppresses false matches in the ventral visual cortex. Another possibility is that the increased excitation during anticorrelated disparity presentation relates to perception of number of dots. Indeed, anticorrelated stimuli can sometimes appear like they contain more dots, although the number of dots is constant between correlated and anticorrelated. This can be explained by a failure to match left and right eye dots during anticorrelation, which can cause a disordered state compared to the binocularly fused percept during binocular correlation (Julesz & Tyler, 1976). In anticorrelated RDS, no coherent object is perceived and therefore LO responds to the individual dots, possibly causing greater excitation. The ventral visual cortex is highly sensitive to numerosity(Cai et al., 2023; Paul et al., 2022), and an increase in cortical excitability could be due to the greater number of dots processed in LO.

### Evidence for robust E/I-balance in ventral visual cortex

The balance between excitation and inhibition in the early visual cortex regulates neural processing and plasticity (Hensch, 2005). We assessed the balance between Glx and GABA+ in the early and ventral visual cortex by correlating the concentrations across participants. Initially, Glx and GABA+ correlated in both the early visual cortex and ventral visual cortex. However, controlling for confounding factors, including spectral quality, age, sex and model fit error demonstrated that the correlation was robust in the ventral but not early visual cortex. Prior studies have shown region-specificity of E/I balance, some reporting a balance in the medial parietal cortex (Steel et al., 2020), and others reporting no balance in the visual and motor cortex(Rideaux, 2021). However, a subsequent study by Rideaux et al., (Rideaux et al., 2022) demonstrated E/I balance in the early visual cortex and the discrepancy was suggested to originate from the proxy of excitation used. While using Glutamate+Glutamine at 3T may be less reliable than Glutamate at 7T (Rideaux 2022), our results using the 3T measurement support the existence of E/I balance in the ventral visual cortex of the human brain. This fine balance between excitation and inhibition may increase sensitivity to objects in a manner that is robust to noise (Rubin et al., 2017). Selectivity to complex objects, like faces, emerges gradually over the course of development (Grill-Spector et al., 2008).

### Object selective response correlates with Glx during anticorrelated disparity in LO

We found a voxel- and condition-specific correlation between object-selective response and Glx in LO. Our results suggest that individuals who have a higher BOLD-response to objects also have more Glx in area LO, specifically during viewing of anticorrelated binocular disparity (**Fig. 7c**). This correlation was specific to LO, robust to controlling for confounding factors including spectral quality, general excitability and percentage voxel overlap. It is possible that increased Glx during anticorrelated condition reflects adaptation to the stimulus that would have caused an initial inhibitory response. This hypothesis remains untested in our study but would be consistent with a prior EEG study (Rideaux et al., 2020). Although the stimuli were different in appearance and dimension, greater sensitivity in LO to objects, in general, could explain higher Glx to suppress irrelevant disparity cues, i.e. anticorrelated stimuli, and a stronger response objects. Regressing out EVC anticorrelated Glx concentration values did not weaken the correlation, suggesting that it was not driven by general excitability. This result further points towards excitation in LO as a valuable way to investigate disparity processing.

## Limitations

Our study recruited 18 participants, yet some conditions could not be collected from all participants. In addition, quality control reduced the sample size further, with a minimum of 15 participants per condition. Our study did not assess depth perception using a behavioural task. This means that the relevance of the Glx and GABA+ changes cannot be directly related to depth perception.

## Conclusion

We measured the concentration of GABA+ and Glx in the early and ventral visual cortex and demonstrated changes in neurochemistry with correlated and anticorrelated binocular disparity. Our results are consistent with previous work suggesting that cortical neurotransmission is involved in suppressing anticorrelated disparity. However, the direction and type of metabolite were not consistent with prior literature. Unlike previously suggested, we found no evidence that GABA inhibited anticorrelated disparity. Instead, we found a decrease in excitation in EVC, suggesting that glutamatergic neurotransmission and metabolism may be involved. In LO, we found an increase in excitation with anticorrelation, possibly reflecting adaptation to prolonged inhibition or more perceived dots in the absence of a coherent percept.

## Acknowledgements

The authors would like to thank the radiographers for their help in collecting data and the volunteers who took part in the study.

## References

Abdolrahmani, M., Doi, T., Shiozaki, H. M., & Fujita, I. (2016). Pooled, but not single-neuron, responses in macaque V4 represent a solution to the stereo correspondence problem. Journal of Neurophysiology, 115(4), 1917–1931. 10.1152/jn.00487.2015

Alvarez, I., Hurley, S. A., Parker, A. J., & Bridge, H. (2021). Human primary visual cortex shows larger population receptive fields for binocular disparity-defined stimuli. Brain Struct Funct, 226(9), 2819–2838. 10.1007/s00429-021-02351-3

Bates D M. M., Bolker B, Walker S. (2015). Fitting Linear Mixed-Effects Models Using lme4. Journal of Statistical Software, 67(1), 1–48. 10.18637/jss.v067.i01

Bednarik, P., Tkac, I., Giove, F., DiNuzzo, M., Deelchand, D. K., Emir, U. E., Eberly, L. E., & Mangia, S. (2015). Neurochemical and BOLD responses during neuronal activation measured in the human visual cortex at 7 Tesla. J Cereb Blood Flow Metab, 35(4), 601–610. 10.1038/jcbfm.2014.233

Bridge, H., & Parker, A. J. (2007). Topographical representation of binocular depth in the human visual cortex using fMRI. J Vis, 7(14), 15 11–14. 10.1167/7.14.15

Cai, Y., Hofstetter, S., & Dumoulin, S. O. (2023). Nonsymbolic Numerosity Maps at the Occipitotemporal Cortex Respond to Symbolic Numbers. Journal of Neuroscience, 43(16), 2950–2959. 10.1523/JNEUROSCI.0687-22.2023

Clarke, W. T., Bell, T. K., Emir, U. E., Mikkelsen, M., Oeltzschner, G., Shamaei, A., Soher, B. J., & Wilson, M. (2022). NIfTI-MRS: A standard data format for magnetic resonance spectroscopy. Magn Reson Med, 88(6), 2358–2370. 10.1002/mrm.29418

Clarke, W. T., Stagg, C. J., & Jbabdi, S. (2021). FSL-MRS: An end-to-end spectroscopy analysis package. Magn Reson Med, 85(6), 2950–2964. 10.1002/mrm.28630

Cogan, A. I., Lomakin, A. J., & Rossi, A. F. (1993). Depth in anticorrelated stereograms: effects of spatial density and interocular delay. Vision Res, 33(14), 1959–1975. 10.1016/0042-6989(93)90021-n

Cumming, B. G., & Parker, A. J. (1997). Responses of primary visual cortical neurons to binocular disparity without depth perception. Nature, 389(6648), 280–283. 10.1038/38487

Cumming, B. G., Shapiro, S. E., & Parker, A. J. (1998). Disparity detection in anticorrelated stereograms. Perception, 27(11), 1367–1377. 10.1068/p271367

Decramer, T., Premereur, E., Uytterhoeven, M., Van Paesschen, W., van Loon, J., Janssen, P., & Theys, T. (2019). Single-cell selectivity and functional architecture of human lateral occipital complex (vol 17, e3000280, 2019). Plos Biology, 17(12). 10.1371/journal.pbio.3000280

Grill-Spector, K., Golarai, G., & Gabrieli, J. (2008). Developmental neuroimaging of the human ventral visual cortex. Trends Cogn Sci, 12(4), 152–162. 10.1016/j.tics.2008.01.009

Hensch, T. K. (2005). Critical period plasticity in local cortical circuits. Nat Rev Neurosci, 6(11), 877–888. 10.1038/nrn1787

Ip, I. B., Alvarez, I., Tacon, M., Parker, A. J., & Bridge, H. (2022). MRI Stereoscope: A Miniature Stereoscope for Human Neuroimaging. Eneuro, 9(1). 10.1523/ENEURO.0382-21.2021

Ip, I. B., Berrington, A., Hess, A. T., Parker, A. J., Emir, U. E., & Bridge, H. (2017). Combined fMRI-MRS acquires simultaneous glutamate and BOLD-fMRI signals in the human brain. Neuroimage, 155, 113–119. 10.1016/j.neuroimage.2017.04.030

Ip, I. B., Minini, L., Dow, J., Parker, A. J., & Bridge, H. (2014). Responses to interocular disparity correlation in the human cerebral cortex. Ophthalmic and Physiological Optics, 34(2), 186–198. 10.1111/opo.12121

Janssen, P., Vogels, R., Liu, Y., & Orban, G. A. (2003). At least at the level of inferior temporal cortex, the stereo correspondence problem is solved. Neuron, 37(4), 693–701. 10.1016/s0896-6273(03)00023-0

Janssen, P., Vogels, R., & Orban, C. A. (2000a). Selectivity for 3D shape that reveals distinct areas within macaque inferior temporal cortex. Science, 288(5473), 2054–+. 10.1126/science.288.5473.2054

Janssen, P., Vogels, R., & Orban, G. A. (2000b). Three-dimensional shape coding in inferior temporal cortex. Neuron, 27(2), 385–397. 10.1016/s0896-6273(00)00045-3

Julesz, B., & Tyler, C. W. (1976). Neurontropy, an entropy-like measure of neural correlation, in binocular fusion and rivalry. Biol Cybern, 23(1), 25–32. 10.1007/BF00344148

Kourtzi, Z., & Kanwisher, N. (2000). Cortical regions involved in perceiving object shape. Journal of Neuroscience, 20(9), 3310–3318. 10.1523/JNEUROSCI.20-09-03310.2000

Krug, K., Cumming, B. G., & Parker, A. J. (2004). Comparing perceptual signals of single V5/MT neurons in two binocular depth tasks. Journal of Neurophysiology, 92(3), 1586–1596. 10.1152/jn.00851.2003

Kumano, H., Tanabe, S., & Fujita, I. (2008). Spatial frequency integration for binocular correspondence in macaque area V4. Journal of Neurophysiology, 99(1), 402–408. 10.1152/jn.00096.2007

Kurcyus, K., Annac, E., Hanning, N. M., Harris, A. D., Oeltzschner, G., Edden, R., & Riedl, V. (2018). Opposite Dynamics of GABA and Glutamate Levels in the Occipital Cortex during Visual Processing. Journal of Neuroscience, 38(46), 9967–9976. 10.1523/JNEUROSCI.1214-18.2018

Lohia, K., Soans, R. S., Saxena, R., Mahajan, K., & Gandhi, T. K. (2024). Distinct rich and diverse clubs regulate coarse and fine binocular disparity processing: Evidence from stereoscopic task-based fMRI. iScience, 27(6), 109831. 10.1016/j.isci.2024.109831

Mangia, S., Tkac, I., Gruetter, R., Van De Moortele, P. F., Giove, F., Maraviglia, B., & Ugurbil, K. (2006). Sensitivity of single-voxel 1H-MRS in investigating the metabolism of the activated human visual cortex at 7 T. Magn Reson Imaging, 24(4), 343–348. 10.1016/j.mri.2005.12.023

Marr, D., & Poggio, T. (1979). A computational theory of human stereo vision. Proc R Soc Lond B Biol Sci, 204(1156), 301–328. 10.1098/rspb.1979.0029

Paul, J. M., van Ackooij, M., Ten Cate, T. C., & Harvey, B. M. (2022). Numerosity tuning in human association cortices and local image contrast representations in early visual cortex. Nat Commun, 13(1), 1340. 10.1038/s41467-022-29030-z

Preston, T. J., Li, S., Kourtzi, Z., & Welchman, A. E. (2008). Multivoxel Pattern Selectivity for Perceptually Relevant Binocular Disparities in the Human Brain. Journal of Neuroscience, 28(44), 11315–11327. 10.1523/Jneurosci.2728-08.2008

Read, J. C., & Cumming, B. G. (2007). Sensors for impossible stimuli may solve the stereo correspondence problem. Nat Neurosci, 10(10), 1322–1328. 10.1038/nn1951

Read, J. C., Parker, A. J., & Cumming, B. G. (2002). A simple model accounts for the response of disparity-tuned V1 neurons to anticorrelated images. Vis Neurosci, 19(6), 735–753. 10.1017/s0952523802196052

Rideaux, R. (2021). No balance between glutamate+glutamine and GABA+ in visual or motor cortices of the human brain: A magnetic resonance spectroscopy study. Neuroimage, 237, 118191. 10.1016/j.neuroimage.2021.118191

Rideaux, R., Ehrhardt, S. E., Wards, Y., Filmer, H. L., Jin, J., Deelchand, D. K., Marjanska, M., Mattingley, J. B., & Dux, P. E. (2022). On the relationship between GABA+ and glutamate across the brain. Neuroimage, 257, 119273. 10.1016/j.neuroimage.2022.119273

Rideaux, R., Michael, E., & Welchman, A. E. (2020). Adaptation to Binocular Anticorrelation Results in Increased Neural Excitability. J Cogn Neurosci, 32(1), 100–110. 10.1162/jocn_a_01471

Rose, D., Bradshaw, M. F., & Hibbard, P. B. (2003). Attention affects the stereoscopic depth aftereffect. Perception, 32(5), 635–640. 10.1068/p3324

Rothman, D. L., Behar, K. L., Prichard, J. W., & Petroff, O. A. (1997). Homocarnosine and the measurement of neuronal pH in patients with epilepsy. Magn Reson Med, 38(6), 924–929. 10.1002/mrm.1910380611

Rubin, R., Abbott, L. F., & Sompolinsky, H. (2017). Balanced excitation and inhibition are required for high-capacity, noise-robust neuronal selectivity. Proc Natl Acad Sci U S A, 114(44), E9366–E9375. 10.1073/pnas.1705841114

Samonds, J. M., Potetz, B. R., Tyler, C. W., & Lee, T. S. (2013). Recurrent connectivity can account for the dynamics of disparity processing in V1. Journal of Neuroscience, 33(7), 2934–2946. 10.1523/JNEUROSCI.2952-12.2013

Steel, A., Mikkelsen, M., Edden, R. A. E., & Robertson, C. E. (2020). Regional balance between glutamate plus glutamine and GABA plus in the resting human brain. Neuroimage, 220. 10.1016/j.neuroimage.2020.117112

Takemura, A., Inoue, Y., Kawano, K., Quaia, C., & Miles, F. A. (2001). Single-unit activity in cortical area MST associated with disparity-vergence eye movements: evidence for population coding. Journal of Neurophysiology, 85(5), 2245–2266. 10.1152/jn.2001.85.5.2245

Tanabe, S., & Cumming, B. G. (2014). Delayed suppression shapes disparity selective responses in monkey V1. Journal of Neurophysiology, 111(9), 1759–1769. 10.1152/jn.00426.2013

Tanabe, S., Haefner, R. M., & Cumming, B. G. (2011). Suppressive mechanisms in monkey V1 help to solve the stereo correspondence problem. Journal of Neuroscience, 31(22), 8295–8305. 10.1523/JNEUROSCI.5000-10.2011

Tanabe, S., Umeda, K., & Fujita, I. (2004). Rejection of false matches for binocular correspondence in macaque visual cortical area V4. Journal of Neuroscience, 24(37), 8170–8180. 10.1523/JNEUROSCI.5292-03.2004

Tanabe, S., Yasuoka, S., & Fujita, I. (2008). Disparity-energy signals in perceived stereoscopic depth. J Vis, 8(3), 22 21–10. 10.1167/8.3.22

Vallat, R. (2018). Pingouin: statistics in Python. Journal of Open Source Software, 3(31), 1026. 10.21105/joss.01026

Verhoef, B. E., Vogels, R., & Janssen, P. (2016). Binocular depth processing in the ventral visual pathway. Philos Trans R Soc Lond B Biol Sci, 371(1697). 10.1098/rstb.2015.0259

Willis, H. E., Ip, I. B., Watt, A., Campbell, J., Jbabdi, S., Clarke, W. T., Cavanaugh, M. R., Huxlin, K. R., Watkins, K. E., Tamietto, M., & Bridge, H. (2023). GABA and Glutamate in hMT+ Link to Individual Differences in Residual Visual Function After Occipital Stroke. Stroke, 54(9), 2286–2295. 10.1161/STROKEAHA.123.043269

Yoshioka, T. W., Doi, T., Abdolrahmani, M., & Fujita, I. (2021). Specialized contributions of mid-tier stages of dorsal and ventral pathways to stereoscopic processing in macaque. Elife, 10. 10.7554/eLife.58749

Zou, B., Liu, Y., & Wolfe, J. M. (2022). Top-down control of attention by stereoscopic depth. Vision Res, 198, 108061. 10.1016/j.visres.2022.108061

