## Supplementary Materials for "Correlated and Anticorrelated Binocular Disparity Modulate GABA+ and Glutamate/glutamine Concentrations in the Human Visual Cortex"

| Condition | EVC GABA+ | Glx | LO GABA+ | Glx |
|---|---|---|---|---|
| **Rest** | 17 | 17 | 16 | 15 |
| **Anti** | 16 | 17 | 18 | 16 |
| **Corr** | 16 | 15 | 17 | 16 |

**Table S1**. Total number of data points for each condition and voxel.

| Voxel | Rest FWHM (Hz) | Corr FWHM (Hz) | Anti FWHM (Hz) | Rest abs CRLB | Corr abs CRLB | Anti abs CRLB |
|---|---|---|---|---|---|---|
| **EVC** | 5.37 ± 0.47 | 5.31 ± 0.52 | 5.32 ± 0.49 | 0.21 ± 0.03 | 0.21 ± 0.03 | 0.21 ± 0.02 |
| **LO** | 6.11 ± 0.49 | 6.04 ± 0.53 | 6.06 ± 0.51 | 0.21 ± 0.03 | 0.21 ± 0.02 | 0.21 ± 0.03 |

**Table S2.** Measures of spectral quality and GABA+ fit error. Mean ±standard deviation.

| Pt | Stimulus Type | Max | Peak disparity over time in degrees |
|---|---|---|---|
| 01 | Squares | ±0.3 | 0.0500, 0.0517, 0.0545, 0.0592, 0.0670, 0.0801, 0.1019, 0.1382, 0.1989, 0.3000 |
| 02 | Squares | ±0.3 | 0.0500, 0.0517, 0.0545, 0.0592, 0.0670, 0.0801, 0.1019, 0.1382, 0.1989, 0.3000 |
| 03 | Squares | ±0.3 | 0.0500, 0.0517, 0.0545, 0.0592, 0.0670, 0.0801, 0.1019, 0.1382, 0.1989, 0.3000 |
| 04 | Squares | ±0.3 | 0.0500, 0.0517, 0.0545, 0.0592, 0.0670, 0.0801, 0.1019, 0.1382, 0.1989, 0.3000 |
| 05 | Squares | ±0.3 | 0.0500, 0.0517, 0.0545, 0.0592, 0.0670, 0.0801, 0.1019, 0.1382, 0.1989, 0.3000 |
| 06* | Squares | ±0.3 | 0.0500, 0.0517, 0.0545, 0.0592, 0.0670, 0.0801, 0.1019, 0.1382, 0.1989, 0.3000 |
| 07 | Sinusoid | ±0.6 | 0.1000, 0.1034, 0.1090, 0.1184, 0.1340, 0.1602, 0.2038, 0.2764, 0.3978, 0.6000 |
| 08 | Sinusoid | ±0.6 | 0.1000, 0.1034, 0.1090, 0.1184, 0.1340, 0.1602, 0.2038, 0.2764, 0.3978, 0.6000 |
| 09 | Sinusoid | ±0.6 | 0.1000, 0.1034, 0.1090, 0.1184, 0.1340, 0.1602, 0.2038, 0.2764, 0.3978, 0.6000 |
| 10 | Sinusoid | ±0.6 | 0.1000, 0.1034, 0.1090, 0.1184, 0.1340, 0.1602, 0.2038, 0.2764, 0.3978, 0.6000 |
| 11 | Sinusoid | ±0.6 | 0.1000, 0.1034, 0.1090, 0.1184, 0.1340, 0.1602, 0.2038, 0.2764, 0.3978, 0.6000 |
| 12 | Sinusoid | ±0.6 | 0.1000, 0.1034, 0.1090, 0.1184, 0.1340, 0.1602, 0.2038, 0.2764, 0.3978, 0.6000 |
| 13 | Sinusoid | ±0.6 | 0, 0.1000, 0.1034, 0.1090, 0.1184, 0.1340, 0.1602, 0.2038, 0.2764, 0.3978, 0.6000 |
| 14 | Sinusoid | ±0.6 | 0, 0.1000, 0.1034, 0.1090, 0.1184, 0.1340, 0.1602, 0.2038, 0.2764, 0.3978, 0.6000 |
| 15 | Sinusoid | ±0.6 | 0, 0.1000, 0.1034, 0.1090, 0.1184, 0.1340, 0.1602, 0.2038, 0.2764, 0.3978, 0.6000 |
| 16 | Sinusoid | ±0.6 | 0, 0.1000, 0.1034, 0.1090, 0.1184, 0.1340, 0.1602, 0.2038, 0.2764, 0.3978, 0.6000 |
| 17 | Sinusoid | ±0.6 | 0, 0.1000, 0.1034, 0.1090, 0.1184, 0.1340, 0.1602, 0.2038, 0.2764, 0.3978, 0.6000 |
| 18 | Sinusoid | ±0.6 | 0, 0.1000, 0.1034, 0.1090, 0.1184, 0.1340, 0.1602, 0.2038, 0.2764, 0.3978, 0.6000 |

**Table S3:** Random dot stereograms by participant. Columns report stimulus type, maximum disparity, and individual disparity steps within a cycle. *data imputed.

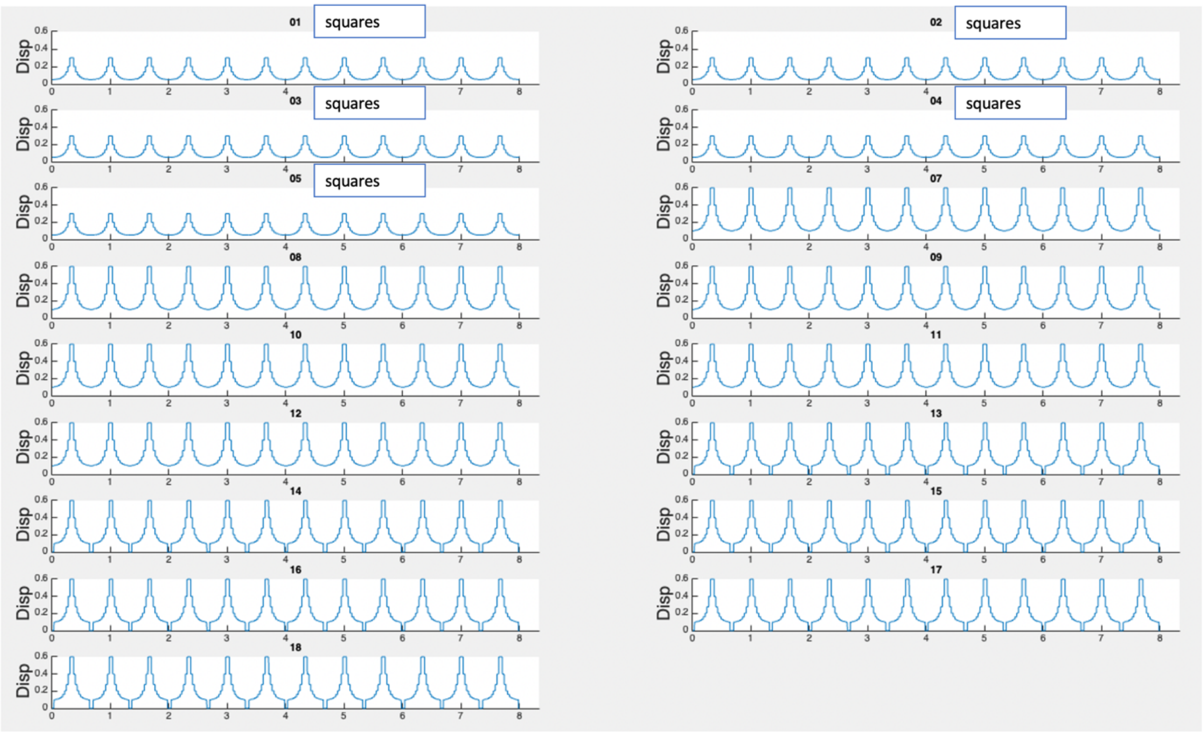


Fig S1: Absolute disparity modulation over 8 min runtime. Y-axis shows absolute disparity. X-axis shows time in minutes.

**Table S3:** Minimum Reporting Standards for MRS Checklist

| Site (Name or Number)<br><br>1. Hardware | 1 |
|---|---|
| a. Field strength [T] | 3T |
| b. Manufacturer | Siemens |
| c. Model (software version if available) | Prisma (VE11E) |
| d. RF coils: nuclei (transmit/ receive), number of channels, type, body part | Siemens 64-channel head coil. |
| e. Additional hardware | N/A |
| **2. Acquisition** | |
| a. Pulse sequence | MEGA-PRESS<br><br>We used a locally developed MEGA-PRESS sequence, derived from the CMRR spectroscopy package MEGA-PRESS sequence (https://www.cmrr.umn.edu/spectro/).<br><br>1. Tremblay S, Beaulé V, Proulx S, Lafleur LP, Doyon J, Marjańska M, Théoret H. The use of magnetic resonance spectroscopy as a tool for the measurement of bi- |

| | |
|---|---|
| | hemispheric transcranial electric stimulation effects on primary motor cortex metabolism. J Vis Exp 2014;93,e51631.<br>2. Marjańska M, Lehéricy S, Valabrègue R, Popa T, Worbe Y, Russo M, Auerbach EJ, Grabli D, Bonnet C, Gallea C, Coudert M, Yahia-Cherif L, Vidailhet M, Meunier C. Brain dynamic neurochemical changes in dystonic patients: a magnetic resonance spectroscopy study. Mov Disord. 2013;28:201-9. |
| b. Volume of Interest (VOI) locations | Primary visual cortex voxel (EVC)<br><br>Lateral occipital cortex (LO) |
| c. Nominal VOI size [$cm^3$, $mm^3$] | Anterior - Posterior: 20 mm<br><br>Left - Right: 25 mm<br><br>Head - Foot: 25 mm |
| d. Repetition Time (TR), Echo Time (TE) [ms, s] | TR = 1500ms<br><br>TE = 68ms |
| e. Total number of Excitations or acquisitions per spectrum<br><br>In time series for kinetic studies<br><br>i. Number of Averaged spectra (NA) per time-point<br>ii. Averaging method (e.g. block-wise or moving average)<br>iii. Total number of spectra (acquired / in time-series) | 160 edit-on and 160 edit-off per editing condition (320 total) |
| f. Additional sequence parameters<br><br>(spectral width in Hz, number of spectral points, frequency offsets)<br><br>If STEAM:, Mixing Time (TM)<br><br>If MRSI: 2D or 3D, FOV in all directions, matrix size, acceleration factors, sampling method | 4000Hz<br><br>2048 points |
| g. Water Suppression Method | VAPOR with additional water suppression using dual-band editing pulse. |
| h. Shimming Method, reference peak, and thresholds for "acceptance of shim" chosen | Automated, vendor supplied 3D GRE $B_0$ field mapping technique ("GRE Brain") was used to ensure that vendor-reported full-width-at-half-maximum (FWHM) were below 20 Hz, and that water-unsuppressed MRS-measured FWHM were <12 Hz. |

| | |
|---|---|
| i. Triggering or motion correction method<br><br>(respiratory, peripheral, cardiac triggering, incl. device used and delays) | None |
| **3. Data analysis methods and outputs** | |
| a. Analysis software | FSL-MRS 2.1.19 |
| b. Processing steps deviating from quoted reference or product | fsl_mrs_preproc_edit \ <br>--data data.nii.gz \ # metabolite file<br>--reference data_phasecorr.nii.gz \ # phase corrected water reference<br>--output /preproc_data \ # output directory<br>--leftshift 3 \ # preprocessing parameter<br>--hlsvd \ # residual water removal<br>--report \ <br>--overwrite \ <br>--align_window_dynamic 16 \ <br>--align_ppm_edit 2.5 3.5 \ # phase/frequency alignment range |
| c. Output measure<br><br>(e.g. absolute concentration, institutional units, ratio) Processing steps deviating from quoted reference or product | Water scaled and tissue fraction corrected absolute concentrations (mMol/kg). |
| d. Quantification references and assumptions, fitting model assumptions | fsl_mrs \ <br>--data diff.nii.gz \ <br>--h2o wref.nii.gz \ # water reference file<br>--tissue_frac tissue_fraction.json \ # tissue fraction file<br>--basis uzay_svs_mpress_68_with_db/diff \ # basis set<br>--metab_groups sysMM \ # grouping<br>--keep GABA GSH Glu Gln NAA NAAG sysMM \\. # fitting<br>--combine Glu Gln GSH\ <br>--combine GABA sysMM \ <br>--combine NAA NAAG \ <br>--internal_ref NAA \ <br>--baseline_order -1 \ <br>--output /fit_output/data \ |

| | |
|---|---|
| | --overwrite \\<br><br>--report \\<br><br>[1] Clarke WT, Stagg CJ, Jbabdi S. FSL-MRS: An end-to-end spectroscopy analysis package. Magnetic Resonance in Medicine 2021;85:2950-2964 doi: 10.1002/mrm.28630. |
| 4. Data Quality | |
| a. Reported variables<br><br>(SNR, Linewidth (with reference peaks)) | FSL-MRS reported linewidth of the inverted NAA peak (FWHM). |
| b. Data exclusion criteria | Univariate outliers: 1.5 interquartile range in metabolite concentrations. |
| c. Quality measures of postprocessing Model fitting (e.g. CRLB, goodness of fit, SD of residual) | FSL-MRS reported absolute Cramer-Rao lower bounds (CRLB). |
| d. Sample Spectrum | Group spectra for all conditions are shown in **Figure 2** in the Main Document. |
